# Mapping the SARS-CoV-2 spike glycoprotein-derived peptidome presented by HLA class II on dendritic cells

**DOI:** 10.1101/2020.08.19.255901

**Authors:** Robert Parker, Thomas Partridge, Catherine Wormald, Rebeca Kawahara, Victoria Stalls, Maria Aggelakopoulou, Jimmy Parker, Rebecca Powell Doherty, Yoanna Ariosa Morejon, Esther Lee, Kevin Saunders, Barton F. Haynes, Priyamvada Acharya, Morten Thaysen-Andersen, Persephone Borrow, Nicola Ternette

## Abstract

Understanding and eliciting protective immune responses to severe acute respiratory syndrome coronavirus 2 (SARS-CoV-2) is an urgent priority. To facilitate these objectives, we have profiled the repertoire of human leukocyte antigen class II (HLA-II)-bound peptides presented by HLA-DR diverse monocyte-derived dendritic cells pulsed with SARS-CoV-2 spike (S) protein. We identify 209 unique HLA-II-bound peptide sequences, many forming nested sets, which map to sites throughout S including glycosylated regions. Comparison of the glycosylation profile of the S protein to that of the HLA-II-bound S peptides revealed substantial trimming of glycan residues on the latter, likely introduced during antigen processing. Our data also highlight the receptor-binding motif in S1 as a HLA-DR-binding peptide-rich region. Results from this study have application in vaccine design, and will aid analysis of CD4+ T cell responses in infected individuals and vaccine recipients.

## INTRODUCTION

Severe acute respiratory syndrome coronavirus 2 (SARS-CoV-2) is a novel beta-coronavirus that first emerged as a human pathogen in the Hubei province of China in late 2019, and is the aetiologic agent of coronavirus disease 2019 (COVID-19). Although SARS-CoV-2 infection is frequently asymptomatic or results in only mild illness, ∼20% of symptomatically infected individuals progress to develop severe pneumonia, acute respiratory distress syndrome and/or sepsis, which can be fatal. By the 24th of July 2020 15,296,926 cases and 628,903 deaths had been reported worldwide^1^. The rapid global spread of SARS-CoV-2 and resulting pandemic have placed tremendous pressure on healthcare services, had a huge societal impact, and profoundly damaged the global economy, prompting an urgent need for vaccines to prevent further spread of infection and avert disease development^1^.

Immune correlates of protection against SARS-CoV-2 infection and progression to severe disease are not yet well-understood, although infection was found to induce at least-short-term protective immunity in a SARS-CoV-2 non-human primate (NHP) infection model, indicating that immune responses are capable of mediating protection^2^. Passively transferred neutralising antibodies (nAbs) protect against SARS-CoV-2 infection in small animal models, and convalescent sera has been shown to be effective in the treatment of severe disease, suggesting the utility of nAb induction by vaccines^3,4,5^. Notably, the four seasonal common cold-causing human coronaviruses and the zoonotic Middle East respiratory syndrome (MERS) and SARS coronaviruses typically elicit poorly-sustained nAb responses, putatively enabling subsequent re-infection^6^. However, somewhat more durable T cell responses are induced, which in animal models can prevent development of severe disease on challenge, providing a rationale for vaccine-mediated induction of T cell as well as nAb responses^7,8,9,10^. More than 150 candidate SARS-CoV-2 vaccines are now in preclinical development or clinical trials^11^.

The SARS-CoV-2 spike (S) glycoprotein (comprised of S1 and S2 subunits) is the primary target of vaccine development efforts. Homotrimers of the transmembrane S protein on the virion surface mediate virion attachment and entry into host cells, making S a key target for nAbs^12^. S is also highly immunogenic for T cells, with many studies suggesting that although infected individuals mount CD4^+^ and CD8^+^ T cell responses to epitopes throughout the viral proteome, S is often at the top of the antigenic hierarchy^13,14,15^. The relative roles of CD4^+^ and CD8^+^ T cells in disease control or pathogenesis and impact of their protein and epitope specificity are unknown; but given the importance of CD4^+^ T cells (particularly CD4^+^ T follicular helper (Tfh) cells) in providing help for antibody responses^16^, and the correlation of memory B cell/nAb responses to S with circulating CD4^+^ Tfh responses in recovered COVID patients^17^, induction of potent Tfh cell responses to the S protein is likely to be crucial for the success of nAb-inducing vaccines.

CD4^+^ T cells are initially activated in response to recognition of specific peptides presented with major histocompatibility complex class II (MHC-II) molecules on professional antigen presenting cells such as dendritic cells (DCs)^18^. The human MHC-II region encodes polymorphic human leukocyte antigen (HLA)-DRA/DRB1, -DRA/DRB3, -DRA/DRB4, - DRA/DRB5, -DPA1/DPB1 and -DQA1/DQB1 molecules, with *HLA-DRA/DRB1* being expressed at the highest and *HLA-DQA1/DQB1* at the lowest levels^19,20^. HLA-II polymorphisms dictate the repertoire of peptides presented for CD4^+^ T cell recognition and shape the response elicited, which can influence the outcome of infection or vaccination^21^. Here, we defined SARS-CoV-2 S-derived peptides presented with diverse HLA-II alleles on DCs to facilitate analysis of pre-existing, post-infection or vaccine-elicited S-specific CD4^+^ T(fh) cell responses and their roles in protection, pathogenesis and prevention of re-infection.

## RESULTS

### Approach for analysis of SARS-CoV-2 S HLA-II presentation by monocyte-derived DCs (MDDCs)

To identify peptides in the SARS-CoV-2 S protein with potential for targeting by CD4^+^ T cell responses, a mass spectrometry-based immunopeptidome profiling approach was employed to define peptides presented by HLA-II on DCs, antigen presenting cells that play a key role in *in vivo* CD4^+^ T cell priming (Figure 1A). MDDCs were generated from 5 HLA-DRB1-heterozygous donors, selected to enable profiling of peptides presented with a total of 9 different HLA-DRB1 alleles (Table 1); these donors also expressed 7 distinct HLA-DPB1 alleles. MDDCs from each donor were pulsed with a recombinant SARS-CoV-2 S protein vaccine immunogen candidate^22^ (produced in a mammalian cell expression system, Figure S2) or a recombinant viral glycoprotein from an unrelated virus that had been produced in the same way (to provide a negative control dataset), and incubated for 18 hours to allow antigen uptake, processing and presentation. Flow cytometry analysis indicated that, as anticipated, the CD11c^+^ MDDC population had robust expression of HLA-I and HLA-DR, expressed high levels of the lectin-type receptors DC-SIGN and DEC-205 and had a relatively immature phenotype, expressing low levels of the DC maturation marker CD83 and moderate levels of the costimulatory molecules CD40, CD80 and CD86. Notably, no difference was observed in the phenotype of SARS-CoV-2 S protein and control protein-pulsed MDDCs, indicating that the S protein had not altered the DC maturation state or HLA expression levels (Figure 1B, Figure S1).

**Table 1.**
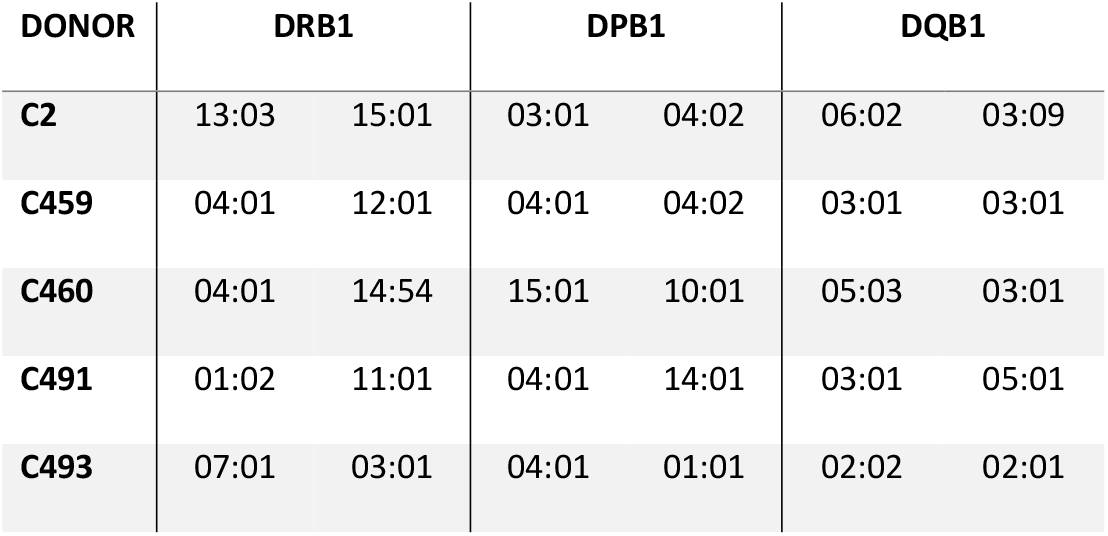
Donor HLA-DRB1, -DPB1 and -DQB1 alleles

| DONOR | DRB1 |  | DPB1 |  | DQB1 |  |
| --- | --- | --- | --- | --- | --- | --- |
| **C2** | 13:03 | 15:01 | 03:01 | 04:02 | 06:02 | 03:09 |
| **C459** | 04:01 | 12:01 | 04:01 | 04:02 | 03:01 | 03:01 |
| **C460** | 04:01 | 14:54 | 15:01 | 10:01 | 05:03 | 03:01 |
| **C491** | 01:02 | 11:01 | 04:01 | 14:01 | 03:01 | 05:01 |
| **C493** | 07:01 | 03:01 | 04:01 | 01:01 | 02:02 | 02:01 |

**Figure 1.**
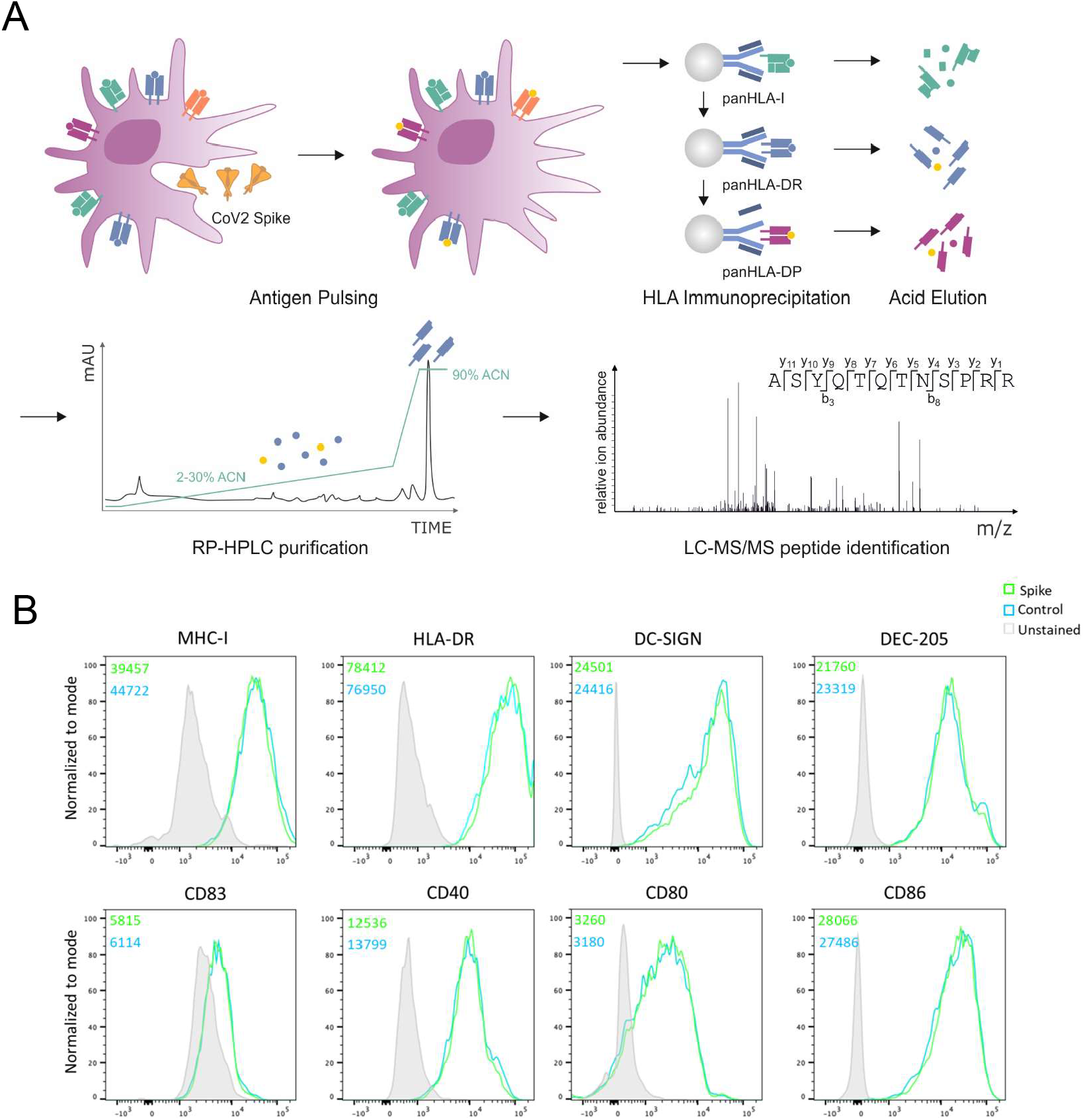
Generation of MDDCs for HLA-II immunopeptidomic profiling. A. Schematic depicting the workflow used to generate the recombinant S protein pulsed MDDC immunopeptidome. MDDCs were pulsed with S or control protein for 18h, and sequential HLA IPs were carried out as indicated. Peptides were eluted from HLA molecules, purified by preparative HPLC and subjected to a LC-MS/MS identification workflow. B. MDDCs were harvested following overnight protein pulse and stained with antibodies to identify DCs and assess their expression of HLA-ABC, HLA-DR, DC-SIGN, DEC-205, CD83, CD40, CD80 and CD86, then analysed by flow cytometry. Histogram plots are gated on live CD11c+CD4+ DCs and show expression of each indicated marker on cells treated with S or control protein. Unstained cells are shown in filled grey histograms. Inset numbers indicate mean fluorescence intensity for each sample.

### Immunopeptidomic profiling of HLA-II-associated peptides presented by MDDCs

Protein-pulsed MDDCs were lysed and sequential immunoprecipitations performed with a pan-HLA-I-specific antibody (W6/32) for depletion of HLA-I complexes, followed by serial pan-HLA-DR (L243) and pan-HLA-DP (B721) immunoprecipitations for enrichment of HLA-DR- and HLA-DP-peptide complexes. After peptide elution and sequencing by tandem mass spectrometry, a total of 27,081 unique HLA-DR- and 2,801 HLA-DP-associated peptide sequences were identified at 1% FDR, of which 147 (HLA-DR) and 12 (HLA-DP) mapped to the S protein (Figure 2A-D, Table S1). None of these peptides were identified in the control protein-pulsed MDDC samples (not shown), consistent with derivation from the S protein antigen. The total number of identified peptides varied in each donor, and was influenced by starting cell numbers (Figure 2A-E). The overall peptide length distributions were highly characteristic of HLA-II-associated peptides, with a median amino acid length of 15 for both human and S peptides (Figure 2F,G). As the HLA-DRA chain is invariant, differences in the peptide binding repertoires of HLA-DR heterodimers are dependent on the HLA-DRB allele expressed. Binding predictions (performed using NetMHCIIpan 4.0) suggested that 60-80% of the peptide sequences identified in the HLA-DR immunopeptidome had a high predicted binding affinity for the donor-specific HLA-DRA/B1 alleles^23^ (Figure 2H). When stratified by HLA-DRB1 allele, ∼80% of peptides were predicted to bind just 1 allele in donors C2, C460 and C459, whilst for donors C491 and C493, peptides were more equally distributed between both HLA-DRB1 alleles (Figure 2I). This was further reflected in an unsupervised Gibbs clustering analysis, which revealed the distinct sequence motifs characteristic of at least one of the donor’s HLA-DRB1 alleles (Figure 2J). Prior to purifying class II complexes, we also purified HLA-I ligands and identified 29,309 self-peptides. There was no evidence of MDDC HLA-I cross-presentation of the pulsed S protein (data not shown).

**Figure 2.**
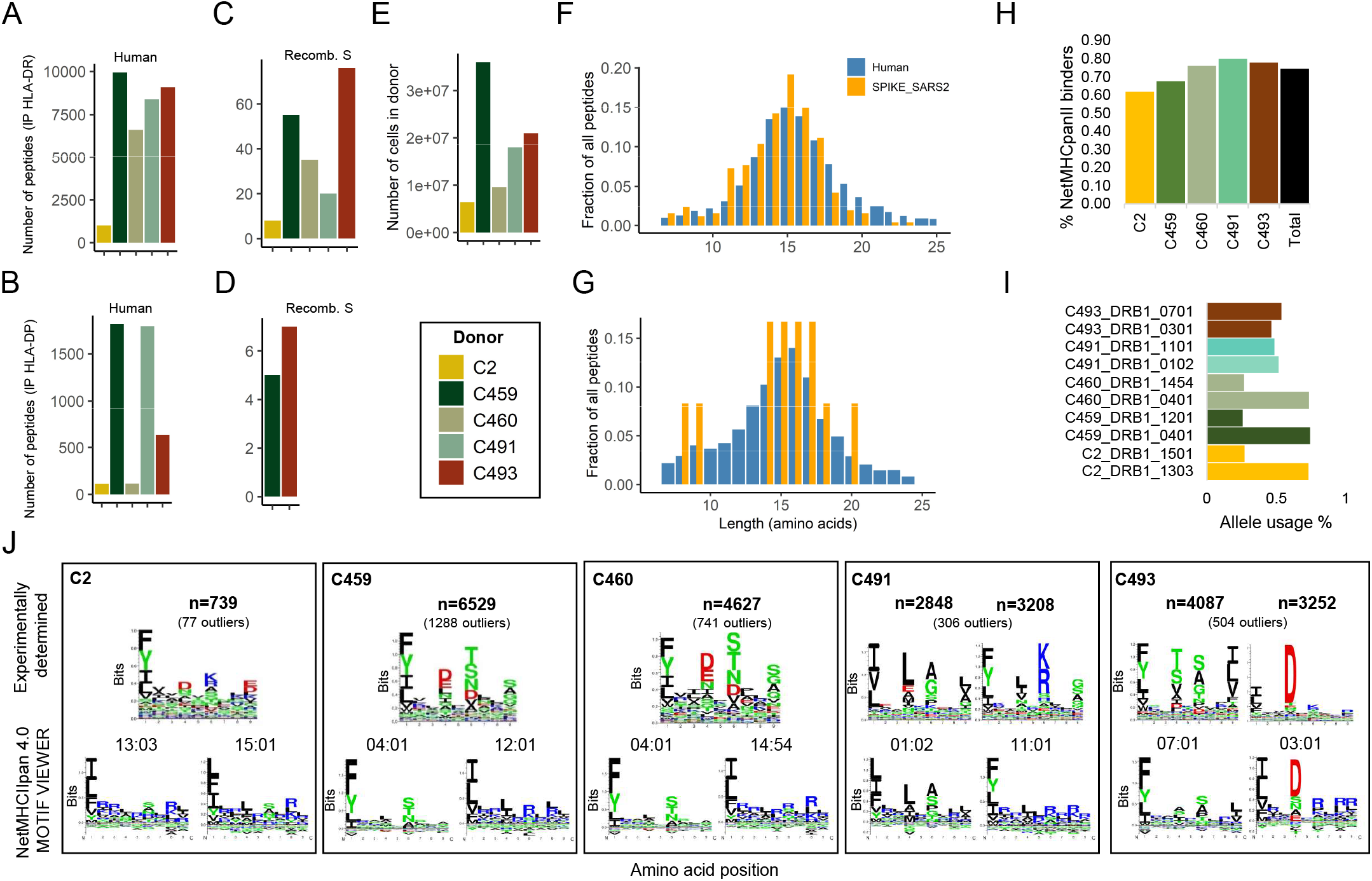
Immunopeptidomic profiling of HLA-DR and HLA-DP bound peptides presented by S-pulsed MDDCs. A-B. Number of unique peptide sequences identified in each HLA-DR (A) or HLA-DP (B) immunoprecipitation sample. C-D. Number of unique peptide sequences that map to S identified in the HLA-DR (C) or HLA-DP (D) samples. E. Total number of MDDCs harvested from each donor. F-G. Length distribution of peptide sequences identified as mapping to human proteins (blue) or S protein (orange). H. Proportion of HLA-DR peptide sequences predicted to bind to the donors’ HLA-DR alleles by NetMHCIIpan 4.0. I. Proportion of predicted HLA-DR binders stratified by allele for each donor. J. All 12-20mers in each sample were clustered using the online (unsupervised) GibbsCluster algorithm. Each cluster is represented by a sequence logo, which corresponds to at least one of the HLA-DRB1 alleles expressed by the donor MDDCs. Amino acids are represented by their single letter code; the more frequently an amino acid occurs a position within peptides, the larger the letter is displayed. The number (n) of peptides within each cluster is indicated along with the number of outlier peptides removed, and clusters are presented with the specific sequence motif for donor DRB1 alleles as reported by NetMHCIIpan 4.0.

### Multiple regions of S are presented by HLA-DR in a genotype-dependent manner

Characteristic of HLA-II-bound peptides, many of the S peptides formed distinctive nested sets around a common core. Two of the identified peptides originated from regions that were altered to assist recombinant protein expression and purification (Figure S2C). The location of the identified S peptides in the context of protein region and domain structure and relative frequency with which particular sites are presented in each donor is summarised in Figure 3A. Several “hotspots” from which a large number of unique peptides (typically different members of a nested set) are presented in multiple donors are apparent across the length of the full S protein, and two regions of S, spanning amino acids 24-49 and 457-485, particularly stand out as the sites from which the highest number of unique HLA-II-bound peptides were derived.

**Figure 3.**
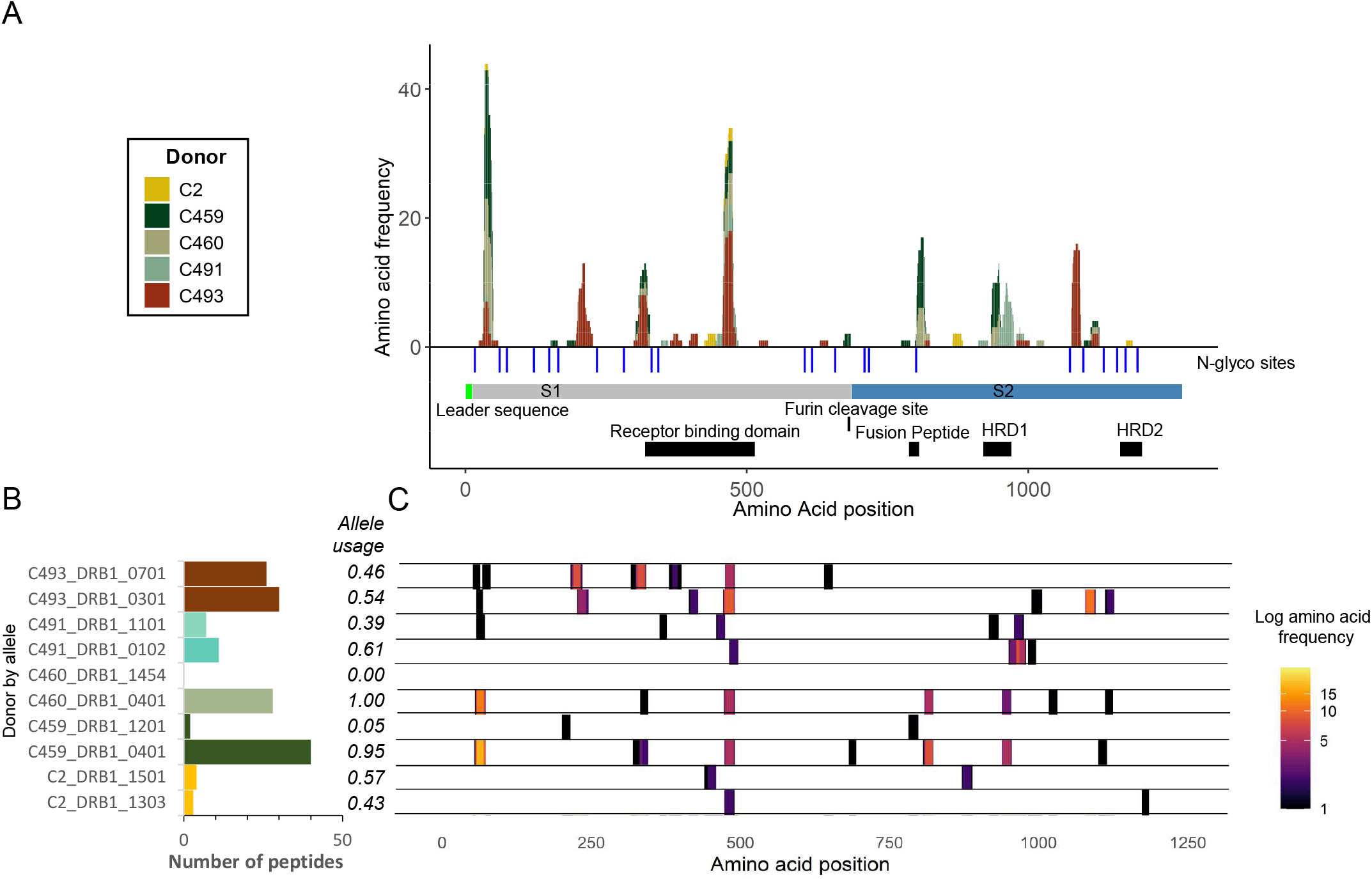
Distribution of HLA-II presented regions within the S glycoprotein. A. Stacked histogram of positional amino acid frequency in S for all peptide sequences identified in the immunopeptidome stratified and coloured by donor. Horizontal bars depict the S1/2 regions, and blue vertical lines indicate N-linked glycosylation sites identified in the purified protein detected by digestion with PNGase F in the presence of heavy water, which leads to a detectable specific mass shift of 2 Da when an N-linked glycan has been removed. The furin cleavage site, receptor binding domain, fusion peptide and heptad repeat domains (HRD) 1 and 2 are depicted as black boxes. B. Number of DR-bound S peptides predicted to bind to the HLA-DRA/B1 alleles expressed in each donor. C. A heatmap of positional amino acid frequency in S for all peptide sequences predicted to bind to donor HLA-DR alleles, stratified by donor and allele on the y axis.

To explore the contribution of individual HLA-DR alleles to presentation of the S protein, we investigated the likely allele to which each peptide was bound using HLA-II binding prediction (NetMHCIIpan 4.0) (Figure 3B,C). 75-100% of the HLA-DR-bound S peptide sequences identified in each donor were predicted to bind one or more of the donor’s HLA-DRB1 alleles (Figure S3A). This stratification demonstrated a within-patient allele usage bias in S presentation that in most donors mirrored the previously observed bias in the percentage of all peptides presented by individual alleles (Figure 3B and Figure 2I). For example, in donor C493 each HLA-DR allele (DRB1*07:01 and DRB1*03:01) had a similar propensity to present both self and S peptides, whilst the majority of peptides were presented with the DRB1*04:01 allele in donor C459 (DRB1*02:01 and DRB1*04:01) and C460 (DRB1*04:01 and DRB1*12:01) MDDCs. A strong correlation was observed between the DRB1*04:01-presented peptide profile of donors C460 and C459, who both expressed the DRB1*04:01 allele (R=0.98), and both donors shared by far the largest number of identical peptide sequences (23) found in any pairwise comparison (Figures S3B,C and S4).

### HLA-II-bound S peptides with N-linked glycosylation predominantly bear truncated paucimannose glycans

To determine the glycosylation status of the S protein immunogen used in this study, a proteomic approach was used to map the N-linked glycosylation sites, involving *in vitro* digestion of the recombinant S protein, trimming of glycans from the generated peptides with PNGaseF in the presence of heavy water (H_2_^18^O), and peptide characterisation by mass spectrometry^24^. This analysis revealed that the 22 N-linked glycosylation sites previously described in S^25^ were occupied in the S protein employed here (Table S2). Notably, regions of the S protein containing glycosites were devoid of peptides identified in our initial analysis of the HLA-II-bound peptidome, raising the question of whether S-derived glycopeptides were also presented by MDDCs (Figure 3A).

To enable glycopeptide analysis, non-PNGaseF-treated S digests were analysed using a well-established glycoproteomics strategy^26^. Using this approach, glycopeptides at 19 sites of S were identified to carry oligomannosidic and complex/hybrid-type *N*-glycans (Figure 4A, Table S3). Most sites displayed extensive glycan microheterogeneity arising from differences in both glycan types and structural features including terminal sialylation and fucosylation in agreement with the known site-specific glycosylation of S^25^.

**Figure 4.**
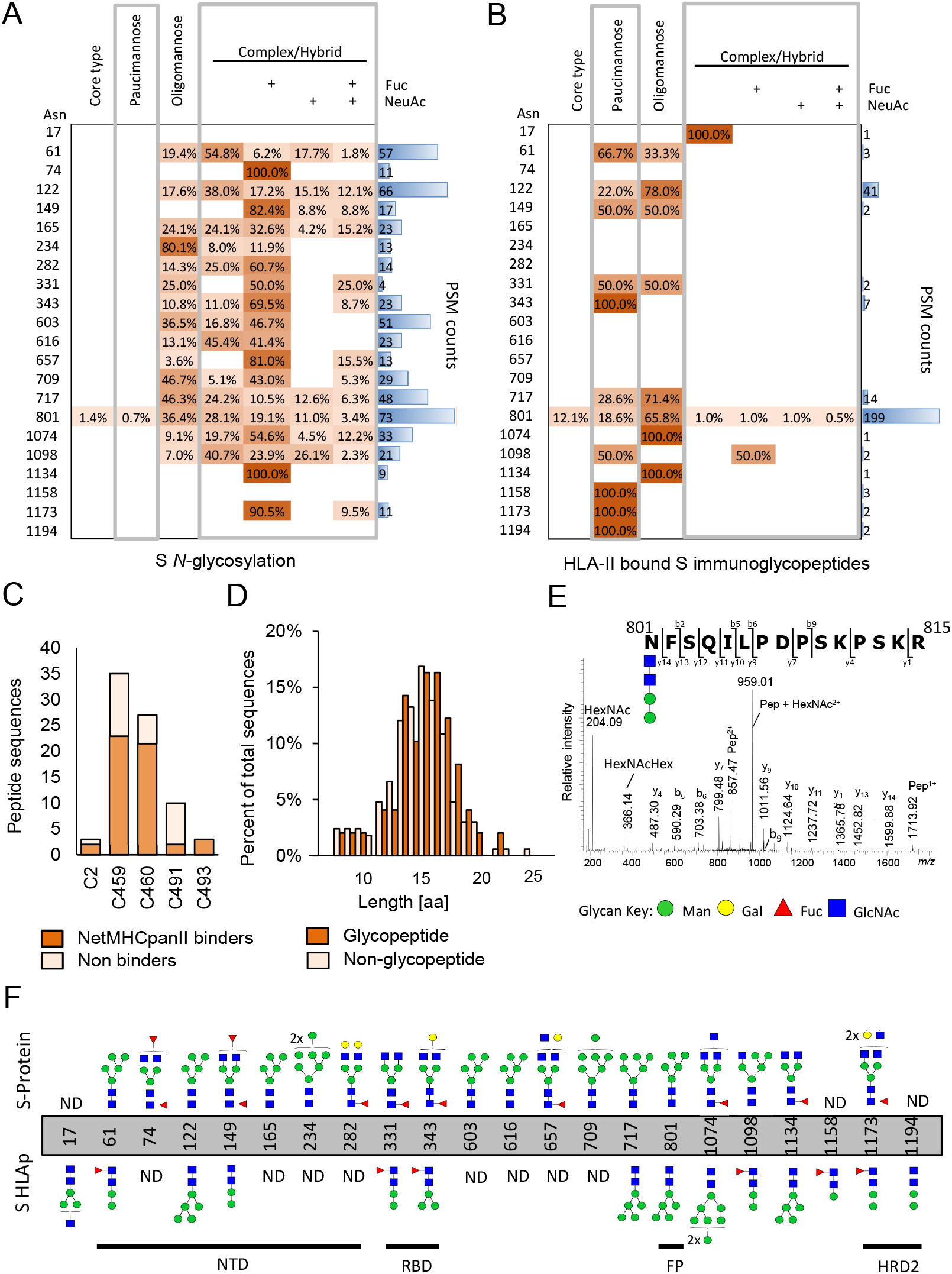
Glycosylation site and glycopeptide analysis of recombinant S protein and HLA-II bound S peptides. A-B. Table showing distribution of glycans identified at each glycosylation site grouped according to the main *N*-glycan types based on intact glycopeptide analysis. A. glycopeptides generated by *in vitro* digestion of the recombinant S protein and B. HLA-II-bound S glycopeptides detected in the immunopeptidome of S-pulsed MDDCs. The heatmap colours indicate the relative frequency of each glycan composition present. The total number of peptide spectral matches (PSM) is reported (blue bars). C. Donor-specific overview of the number of glycosylated peptide sequences and proportion predicted to bind to the donor HLA-DR allele profile. D. Histogram illustrating the amino acid length distribution of glycopeptides and non-modified peptide sequences of S. E. Example of an annotated fragment spectrum of an HLA-II-bound peptide carrying a paucimannosidic-type *N*-glycan in position N801. F. Depiction of the most abundant glycan identified at each *N*-glycosylation sites of the S polypeptide chain prior to MDDC pulsing (top) and of the HLA-II bound S peptides (bottom). The main domains of S are indicated. (Glycan Key: Man = mannose; Gal = galactose; Fuc = fucose; NeuAc = *N*-acetylneuraminic acid; GlcNac = *N*-Acetylglucosamine G).

Next, we applied the site-specific glycopeptide methodology to the mass spectra acquired from samples eluted from HLA-II (Figure 4B). 80 distinct glycopeptide forms mapping to S were identified, the majority of these (76) were derived from the HLA-DR-bound immunopeptidome (Table S4). These glycopeptide forms mapped to 52 unique peptide sequences that typically formed nested sets, were predominantly observed in datasets generated from donors C459 and C460 (where the highest number of unique HLA-DR-bound non-glycosylated peptides were also detected), and had a similar length distribution to S-derived non-glycopeptides (Figure 4C,D). 66% (20-100%) of all glycopeptide sequences were predicted (using NetMHCIIpan 4.0) to bind to one or more of the donor’s HLA-DR alleles (Figure 4C). The largest nested set consisted of glycopeptides from donor C459/C460 MDDCs that mapped to position N801 located directly in the fusion peptide (FP, 788-806), a highly conserved region which facilitates membrane fusion during viral entry (Figure 4E,F). In total, we identified HLA-II-bound glycopeptides bearing glycans derived from 14 of the N-linked glycosylation sites in S (Figure 4F). HLA-II-bound peptides carried predominantly short paucimannosidic-type *N*-glycans while S carried oligomannosidic- and GlcNAc-capped complex-type *N*-glycan structures at these sites (Figure 4B,F). The paucimannosylation of the HLA-II bound peptides comprised both core-fucosylated (M1F, M2F, and M3F i.e. Man1-3GlcNAc2Fuc1) and a fucosylated (M2, Man2GlcNAc2) species as supported by fragment spectra analysis (Figure 4E).

In addition, utilising an open post translational modification (PTM) peptide identification methodology, we identified peptides containing the other most common post translational modifications (other than glycans) in the S immunopeptidome (Figure S5A). The most modified residue was cysteine (C), which was found to have undergone cysteinylation, oxidation to cysteic acid and conversion to glutathione disulphide (Figure S5B). A total of 27 peptides with modified C residues were identified that mapped to 6 positions in S. All peptides contained a single modified C residue and are known to form disulphides in the tertiary structure of S^27^.

### Peptides derived from the receptor binding domain (RBD) of the SARS-CoV-2 S protein are presented by multiple HLA-DR alleles

Altogether, a total of 209 unique HLA-II-bound peptides (differing in amino acid sequence) derived from the SARS-CoV-2 spike protein were detected in this study. The locations and putative presenting HLA-II alleles of these peptides (typically members of large nested sets) are summarised in Figure 5. Partly overlapping nested sets of peptides predicted to be presented by distinct HLA-DR alleles in different donors were identified in several regions of the spike protein. One region in the RBD particularly stood out in this regard, as it contained 5 nested sets of peptides and an additional peptide that were predicted to be presented by 6 different HLA-DR alleles. At least one peptide within this region was found to be presented in every donor studied (Figure 5, Figure 6A). In the donors analysed, a total of 21 unique peptide sequences derived from this region were identified altogether (Figure 6A,B) and versions of some of these post-translationally-modified at residues C480 and C432 also detected (Figure 6C). This region overlapped directly with the receptor binding motif (RBM), an extended insert on the beta-6/5 strands that contains the contact points with the receptor ACE2^28^.

**Figure 5.**
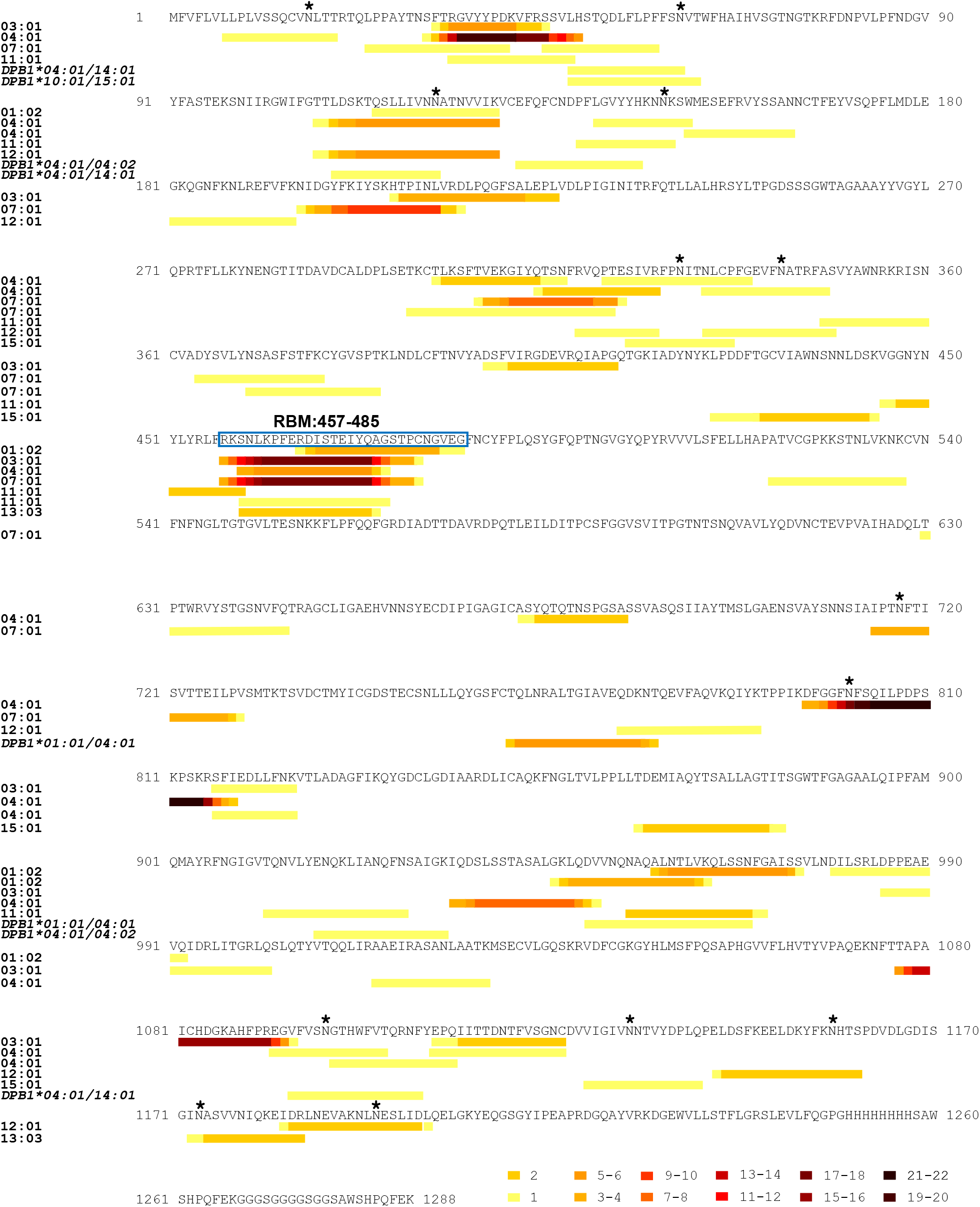
HLA-II peptide mapping across the S protein and RBM. A. Map illustrating the location in the SARS-CoV-2 protein sequence of all the S-derived peptides of 9 to 25 amino acids in length identified in the HLA-II-bound immunopeptidome of S-pulsed MDDCs. HLA-DR-bound peptides are grouped according to the HLA-DR allele to which they had the highest predicted binding affinity (as determined using NetMHCIIpan 4.0), and HLA-DP bound peptides are grouped by donor *HLA-DPB1* type. Within each group, nested peptide sets are indicated in heatmap form, where the colour represents the frequency of each amino acid position within the peptide group. Where a single peptide amino acid sequence was identified multiple times with different sequence modifications the unique amino acid sequence was included only once in positional frequency calculations. Glycosylation sites in S identified in the Byonic analysis of the immunopeptidome are indicated with a * and the nested set covering the RBM is boxed.

**Figure 6.**
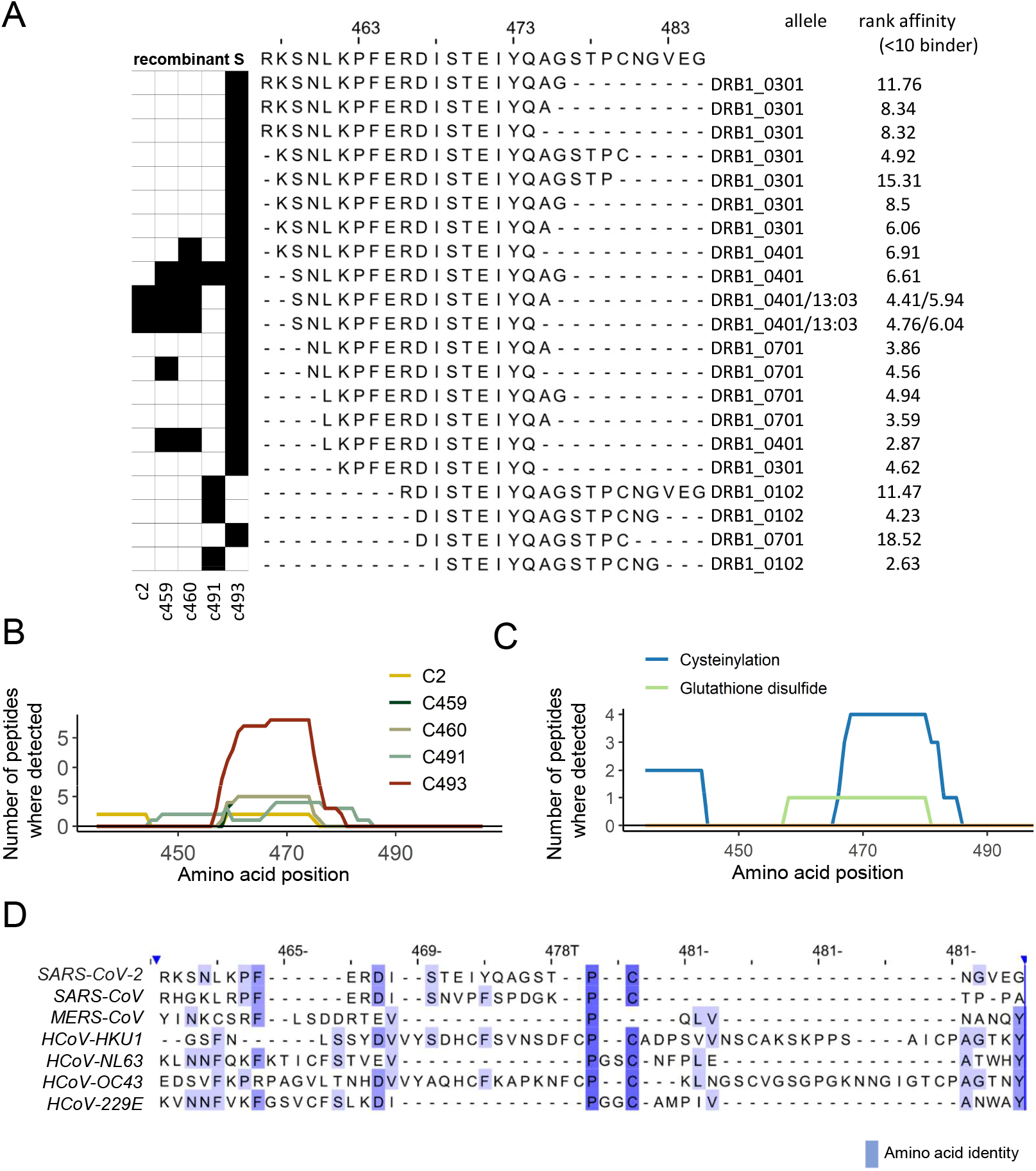
A nested set of unique peptides in the immunopeptidome map to positions 457-485 in the RBD of the full-length S protein. **A**. The black squares on the left indicate the donor(s) in which each peptide was identified. On the right of each peptide sequence, the HLA-DRB1 allele to which it showed the highest predicted affinity of binding (NetMHCIIpan4.0) and rank score is indicated. B. Line plots showing the frequency with which each amino acid position was represented by a peptide identification across the RBD of the S protein in each donor. C. Line plots showing the frequency with which each amino acid position was represented by a cysteine-modified peptide identification (cysteinylated peptides and glutathione-modified peptides) across the RBD of the S protein. D. Alignment of the S protein sequences of the SARS-CoV-2, SARS-CoV, MERS-CoV and 4 endemic human coronaviruses (obtained from Uniprot). Positions relative to the RBD amino acids 457-485 in S are shown. The colour indicates the degree of amino acid identify.

To gain insight into the sequence conservation of this region in other coronaviruses infecting humans, S protein sequences from SARS-CoV-2, SARS-CoV and MERS-CoV (the other beta-coronaviruses that have caused epidemics in humans in the past two decades) and the endemic human coronaviruses 229E, NL63, OC43, and HKU1 were aligned (Figure 6D). Although this region of the SARS-CoV-2 S protein showed some sequence similarity with the equivalent region of the SARS-CoV S protein, this is an indel-rich region of S that was much less well conserved in the other coronaviruses examined. However, although residues that are likely to constitute key anchors in the core regions of the nested peptide sets predicted to bind to particular HLA-DRA/DRB1 molecules (SARS-CoV-2 F464 and S469, which match anchor residue preferences in DRB1*04:01-, 07:01-, 13:03- and 15:01-binding peptides; F464 and D467, which match preferred anchors of DRB1*03:01-binding peptides; and I472 and S477, which match those of DRB1*01:01-binding peptides) are not well conserved or not appropriately positioned relative to one another in all coronaviruses, there appears to be some potential for HLA-II-binding peptides to be generated from this region of other human coronavirus S protein sequences.

## DISCUSSION

Concurrently with the design and clinical evaluation of candidate immunogens in the race to develop vaccines with prophylactic efficacy against SARS-CoV-2 infection and associated disease, there is an urgent need to define T cell epitopes to facilitate analysis of the contribution of T cell responses to protection and pathogenesis in infected individuals and monitoring of immune responses elicited in human vaccine trials^14^. As the SARS-CoV-2 S protein is the major target on the virus for neutralising antibodies^29,3,4,30,5,31, 32, 33^ and has also been shown to be highly immunogenic for T cell responses in infected individuals^13,14,15, 34^, a high proportion of the SARS-CoV-2 vaccines in pre-clinical and clinical development focus on eliciting immune responses to this protein. In this study, we have defined SARS-CoV-2 S-derived peptides presented by DCs (a cell type crucial for induction of immune responses during infection and after vaccination^21^), following uptake and processing of exogenously-acquired S protein. We identify a total of 209 unique HLA-II-bound peptide sequences, including members of 27 nested peptide sets, demonstrating presentation of both glycopeptides and peptides with other post-translational modifications. Notably, our analysis reveals that nested peptide sets derived from a region of the RBD that overlaps with the RBM are presented by diverse HLA-DR alleles, highlighting this as a region of the SARS-CoV-2 S protein that could putatively be targeted by T cell responses in multiple individuals. The peptides identified in this study provide an important resource that will expedite 1) exploration of pre-existing T cell responses to other coronaviruses, 2) cross-comparison of responses elicited by different vaccine immunogens and platforms, and 3) design of next-generation vaccines tailored to elicit enhance responses to nAb epitopes, or focus T cell responses on selected epitopes.

The importance of defining SARS-CoV-2-derived peptides presented by diverse HLA alleles is illustrated by the plethora of recent efforts to employ *in silico* approaches to predict putative T cell epitopes in SARS-CoV-2 proteins^35 36^. Our data give key insight into the repertoire of peptides that are in fact presented with HLA-II when exogenous S protein is internalised and processed by DCs, mimicking a scenario occurring as T cell responses are induced during natural infection or following vaccination with protein immunogens. Whether these peptide profiles are also representative of those presented on DCs in which the S proteins is endogenously-expressed (e.g. as may occur following antigen delivery with viral vectored or nucleic acid-based vaccine platforms), which may lead to antigen processing and peptide association with HLA-II in different intracellular compartments, remains to be determined^18^. Furthermore, although no evidence of *in vitro* cross-presentation of S on HLA-I by MDDCs was observed in this study, vaccine platforms that drive intracellular antigen expression would be expected to result in HLA-I presentation of S-derived peptides, promoting induction of CD8^+^ T cell responses^37,38^.

Our analysis of naturally presented HLA-II-associated peptides from SARS-CoV-2 S protein identified a total of 209 unique HLA-II bound peptides in the 5 donors analysed, including 193 HLA-DR-bound and a further 16 HLA-DP-bound peptides. ∼75% of the HLA-DR bound peptides identified were predicted to bind to at least one of the donor’s HLA-DRB1 alleles, consistent with the higher expression and more dominant antigen presenting role of HLA-DRB1 versus HLA-DRB3, 4 and 5 molecules^17^. Whilst HLA-II binding predictions suggested that the two HLA-DRB1 alleles expressed in some donors made roughly equal contributions to the repertoire of unique peptides presented, in other donors a much greater proportion of the unique peptides detected were predicted to bind to one of their HLA-DRB1 alleles, with DRB1*04:01, a prevalent allele in European populations, appearing to play a more dominant role in antigen presentation in both of the donors expressing this allele. The depth of immunopeptidome profiling achieved for different HLA-DR alleles could be influenced by protein expression levels, but certain alleles may also present a more diverse repertoire of peptides due to a preference for more common amino acids and/or ability to tolerate a greater number of different amino acid residues at key anchor positions, and/or to differences in their association with the peptide editor HLA-DM or the associated HLA-DO protein that modulates its function^18^.

Our analysis identified multiple S-derived peptides in the HLA-II bound repertoire bearing glycans or other post-translational modifications. Viral envelope proteins are often heavily glycosylated and the SARS-CoV2 S protein is no exception^39^, with complex N-glycosylation stemming from 22 sites^25^. Glycosylation of virion surface proteins acts to enhance viral infectivity and also subvert recognitions by host adaptive responses (by shielding nAb binding sites and impairing antigen processing for T cell recognition); but it is also targeted by host innate immune recognition pathways^40^. S protein glycosylation is carried out by the host cell glycan processing machinery, resulting in attachment of a range of oligomannosidic, complex or hybrid structures that mimic mature surface glycoproteins of the host. We initially confirmed that these patterns were present in the intact S protein used to pulse MDDCs. Strikingly, we found that the HLA-II-bound S peptides were in contrast glycosylated at the same site but with glycans rich in highly processed paucimannosidic-type structures. This observation implies a significant modulation of the glycan phenotype upon internalisation, processing, and presentation of the S glycoprotein in MDDCs. Paucimannosidic glycans are defined as truncated α- or β-mannosyl-terminating *N*-glycans carried by proteins expressed widely across the eukaryotic domain, but remains a poorly understood glycan class in human glycobiology and virology^41^. We have recently reported that neutrophils^42, 43^ and monocytes/macrophages^44^, but thus far not DCs, are paucimannose producing cell types in the innate immune system. The paucimannosidic glycans have been proposed to be formed via the sequential trimming facilitated by the *N*-acetyl-β-hexosaminidase isoenzymes and linkage-specific α-mannosidases residing in lysosomes or lysosomal-like compartments^41^. Supporting our data suggesting an extensive DC-driven glycan remodelling ahead of viral glycopeptide presentation, *N*-acetyl-β-hexosaminidase and α-mannosidase, and several other hydrolytic enzymes (e.g. cathepsin D) are known to be abundantly expressed and highly active in MHC class II processing compartments (MIICs)^45^. Furthermore, MHC class II immunopeptides carrying truncated *N*-glycans have previously been reported from other cellular origins^46,47^. CD4+ T cell recognition of glycosylated peptides has been reported in rheumatoid arthritis (O-linked)^48^ and cancer (N-linked)^49^, and CD4^+^ peptides in the melanoma antigen tyrosinase require the presence of N-linked glycosylation to elicit a T cell response^49^. A recent study also showed that immunization of mice with a recombinant human immunodeficiency virus type 1 (HIV-1) envelope (Env) glycoprotein immunogen elicits CD4+ T cell responses to a glycopeptide epitope that provide help for induction of Env-specific antibody responses^50^, suggesting that glycopeptide-targeting CD4+ T cell responses may constitute an important and under-studied component of the immune response elicited following infection or vaccination.

Further investigation of post-translational modifications in HLA-II bound S peptides revealed prevalent cysteine modifications (cysteinylation, glutathione disulphide, cysteine oxidation to cysteic acid). Specifically, we observed cysteinylation and glutathionylation of C479 and C432, a cysteine pair that form two key disulphide bonds in the RBD^28^. Free cysteines are highly reactive and during denaturation can readily become oxidised depending on the surrounding environment^51^. The origin of cysteinylation and glutathionylation in HLA-II peptides is uncertain, and reactions could occur within the endosome or extracellular medium with free cysteine or glutathione, depending on where the peptides are loaded onto MHC molecules^52,^. The existence of cysteine-modified viral epitopes has been explored previously for both class I^53,54^ and class II^55^ epitopes and their presentation in the immunopeptidome is allele- and context-dependent^51^. Biologically, cysteine modifications potentially reflect the redox status of the cell^51^ and can alter T cell recognition of antigens in infection, vaccination and cancer^52, 53, 54, 55^. In agreement with our observations for the SARS-CoV-2 S protein, a CD8^+^ T cell epitope derived from the RBD (312-635) of the S glycoprotein following infection of mice with mouse hepatitis virus (murine coronavirus), in was found to be S-glutathionylated^51^. In cancer, cysteinylation of antigens can confer evasion from T cell recognition; but processing by the IFNy-inducible lysosomal thiol reductase (GILT) can remove antigen cysteinylation and induce antigen processing and T cell responses in the context of melanoma^56^. In an antigen presenting cell loading system similar to that used here, a requirement for peptide endocytosis and processing of a spontaneously cysteinylated peptide was required to establish T cell activation but not MHC binding and presentation^55^. Thus it will be important to determine whether the cysteine-modified S peptides described herein are targeted by T cells following vaccination or during SARS-CoV-2 infection.

Notably, the RBM, an area of the RBD important for interacting with the host receptor ACE2^28^, was found to be a HLA-DR-binding peptide-rich region, with presentation of peptides derived from amino acids 457-485 of the SARS-CoV-2 spike protein being detected in all of the HLA-diverse donors studied. Analysis of the T cell responses elicited following vaccination of mice with recombinant DNA (rDNA) based vectors encoding the S proteins from both SARS and SARS-CoV-2 has shown that the epitopes targeted by CD4+ T cells include a site in the RBD that encompasses the RBM^57,58^, suggesting that peptides derived from this region may be presented in a cross-species manner. Little information is currently available about the epitopes recognised by CD4+ T(fh) cell responses in SARS-CoV-2 infected or vaccinated individuals, although CD4 T cell responses to an epitope at amino acids 449-461 in the SARS-CoV spike (equivalent to, although having a number of sequence differences from amino acids 462-474 of the SARS-CoV-2 spike protein) were detected in healthy donors not exposed to SARS-CoV (or SARS-CoV-2)^36^; and a recent analysis of T cell responses in recovered SARS-CoV-2 infected patients detected a response two overlapping peptides spanning amino acids 446-465 of the SARS-CoV-2 spike protein^59^. More work is needed to determine how commonly this region is recognised by CD4+ T cell responses, both in individuals exposed only to seasonal human coronaviruses and SARS-CoV-2 individuals, and also to explore the inter-virus cross-reactivity of RBM-targeting CD4+ T cells; but our findings highlight this as a putatively immunogenic region worthy of further study.

In summary, our data provide a detailed map of HLA-II-binding peptides in the SARS-CoV-2 S protein that will facilitate the analysis of CD4+ T cell responses to both “conventional” and novel post-translationally-modified epitopes in this important viral target protein. In addition to the utility of our findings in dissection of pre-existing and post-infection responses to the SARS-CoV-2 S protein and their impact on infection outcome, our results also have application in the monitoring of vaccine-elicited immune responses and cross-comparison of the CD4 T(fh) responses elicited by different immunogens, vaccine platforms and immunization regimes. Furthermore, they have important implications for the design of vaccines aiming to target immune responses to specific sites on the S glycoprotein, e.g. indicating the potential for RBD vaccines to elicit CD4+ T(fh) responses in HLA-diverse vaccine recipients, and highlighting regions of the spike protein that may be poor sources of class II epitopes, where linkage to exogenous CD4+ T(fh) epitopes such as broadly-presented peptides from tetanus toxoid may be advantageous in future vaccine design.

## METHODS

### Cell lines and cell culture

Leukocyte cones obtained with appropriate ethical approval from healthy donors who provided written informed consent were purchased from the NHS Blood and Transplant Service, Oxford. All work was compliant with institutional guidelines. Mononuclear cells were isolated from leukocyte cones by separation on a Histopaque-1077 gradient then stored in liquid nitrogen. Genomic DNA was extracted from cells using the QIAamp DNA Mini Kit (QIAGEN), then the HLA-DRB1, -DPB1, and -DQB1 loci were sequenced at the OUH Transplant Laboratory by Sanger sequencing using an ABI-3730 DNA analyzer.

### Protein expression and purification

The SARS-CoV-2 ectodomain constructs were produced and purified as described previously ^22 60^. Briefly, the expression construct included residues 1−1208 of the SARS-CoV-2 S (GenBank: MN908947) with proline substitutions at residues 986 and 987, a “GSAS” substitution at the furin cleavage site (residues 682–685), a C-terminal T4 fibritin trimerization motif, an HRV3C protease cleavage site, a TwinStrepTag and an 8XHisTag. Expression plasmids encoding the ectodomain sequence were used to transiently transfect FreeStyle293F cells using Turbo293 (SpeedBiosystems). Protein was purified on the sixth day post transfection from the filtered supernatant using StrepTactin resin (IBA), followed by size exclusion chromatography using a Superose 6 Increase 10/300 column.

### Differentiation of monocyte-derived DCs (MDDCs)

To differentiate MDDCs, mononuclear cells were thawed and plated at 10^6^ cells per cm^2^ in 20 ml RAB5 (RPMI with 5% human serum AB (Sigma), 2 mM L-alanyl-L-glutamine, 10mM HEPES, 50 units/ml penicillin, 50μg/ml streptomycin) per 175 cm^2^ flask for 2 hours at 37°C. Non-adherent cells were removed by three gentle PBS washes. The remaining adherent cells were cultured in RAB5 containing 300 IU/ml IL-4 and 100 IU/ml GM-CSF for 6 days. MDDCs were harvested on day 6 by incubation in cell dissociation solution (Sigma) followed by gentle scraping. Cells were resuspended in AIM V medium (Thermo Fisher Scientific) and incubated with 0.5 mg of recombinant SARS-CoV-2 S protein or an irrelevant viral envelope glycoprotein produced in the same way for 18 hours. Cells were then harvested by gentle scraping then washed before lysis for HLA immunopurification.

### Flow cytometry

Following differentiation and protein pulse, MDDCs were washed and stained with antibodies specific for HLA-DR (L243, Biolegend), HLA-I (W6/32, Biolegend), DEC-205 (MMRI-7, BD Biosciences), DC-SIGN (9E9A8, Biolegend), CD86 (2331, BD Biosciences), CD83 (HB15e, Biolegend), CD80 (2D10, Biolegend), CD40 (SC3, Biolegend) and CD11c (S-HCL-3, Biolegend) for 20 minutes at room temperature. Cells were then washed and fixed in 4% PFA. Staining data were acquired on a LSRFortessa and analysis was performed on Flowjo software version 10.6.2.

### Enzymatic digestion of SARS-CoV-2 S protein

An equivalent of 5 ug of SARS-CoV-2 S protein was reduced by incubation with 5 mM DTT for 30 minutes. Reduced S was incubated for 30 minutes with 20 mM iodoacetamide (IA) followed by addition of DTT to 20 mM to react with residual IA. 0.2 µg of trypsin or elastase was added per 5 µg of CoV2 S and incubated for 16 hours at 37°C. Sample clean-up was performed with a C18 column (Waters Oasis SPE kit).

### PNGaseF deglycosylation of digested SARS-CoV-2 S protein

Digested CoV2 S protein was dried in a SpeedVac and resuspended in 10 ul 20mM sodium phosphate pH 7.5. Samples were divided equally, dried in a SpeedVac and then resuspended in ^18^O-water (Sigma-Aldrich, 97%) containing 200 U PNGaseF (New England Biolabs) or buffer only. The protein was digested for 16 hours at 37°C. Sample clean-up was performed with a C18 column (Waters Oasis SPE kit) and peptides were dried in a SpeedVac.

### Production and purification of HLA-specific antibodies

Hybridoma cells (clones W6/32, L243, B721) were cultured in a CELLine CL 1000 Bioreactor (Integra) as described in (31). B721 was kindly supplied by Prof. Anthony Purcell, Monash University, Melbourne, Australia. Briefly, cells were cultured in serum-free Hybridoma-SFM medium (Gibco) supplemented with hybridoma mix (2,800 mg/l of d-glucose, 2,300 mg/l of peptone, 2 mM l-glutamine, 1% penicillin/streptomycin, 1% non-essential amino acids, 0.00017% 2-mercaptoethanol). Supernatant containing antibody was harvested and stored at −20. To purify antibodies, supernatants were thawed then centrifuged at 2,500 × g for 25 min at 4°C, filtered (0.2 µm SteriCup Filter (Millipore)) and incubated with protein A resin (PAS) (Expedeon) for 18 hours at 4°C. Antibody-resin complexes were then collected by gravity flow through chromatography columns, washed with 20 ml of PBS, and eluted with 5 ml 100 mM glycine pH 3.0. pH was adjusted to pH 7.4 using 1 M Tris pH 9.5 and antibodies were buffer exchanged into PBS and concentrated with a 5 kDa molecular weight cut-off ultrafiltration device (Millipore).

### Preparation of immunoresin

Cross-linking of purified antibodies was performed as follows: 3 mg of antibody per 0.5 ml PAS was incubated for 1 h at 4°C then washed with 15 ml borate buffer (0.05 M boric acid, 0.05 M KCl, 4 mM NaOH, pH 8.0), 15 ml 0.2 M triethanolamine pH 8.2 and cross-linked to PAS with 15 ml of 40 mM dimethyl pimelimidate in 0.2 M triethanolamine pH 8.3. After 1 h, crosslinking was terminated by addition of 15 ml of 0.2 M Tris pH 8.0, unbound antibody was removed by washing with 15 ml citrate buffer pH 3.0 and three washes with 15 ml PBS.

### HLA complex immunoprecipitation (IP) and peptide purification

Harvest MDDCs were washed in PBS then lysed by mixing for 45 minutes with 3 ml of lysis buffer (0.5% (v/v) IGEPAL 630, 50 mM Tris pH 8.0, 150 mM NaCl and 1 tablet cOmplete Protease Inhibitor Cocktail EDTA-free (Roche) per 10 ml buffer) at 4°C. Lysate clarification was achieved by centrifugation at 3,000 x g for 10 mins followed by a 20,000 × g spin step for 15 mins at 4°C. 3 mg of W6/32 antibody-PAS was incubated with lysate for at least 5 h at 4°C with gentle rotation. Resin was collected by gravity flow and flow-through lysate was sequentially incubated with L243- and then B721-PAS. Antibody-resin-HLA complexes were washed with 15 ml of 0.005% IGEPAL, 50 mM Tris pH 8.0, 150 mM NaCl, 5 mM EDTA, 15 ml of 50 mM Tris pH 8.0, 150 mM NaCl, 15 ml of 50 mM Tris pH 8.0, 450 mM NaCl, and 15 ml of 50 mM Tris pH 8.0. 3 ml of 10% acetic acid was used to elute bound HLA complexes from the PAS-antibody resin, which were then dried by vacuum centrifugation. Eluted peptides were dissolved in loading buffer (0.1% (v/v) trifluoroacetic acid (TFA), 1%(v/v) acetonitrile in water), and then injected by a Ultimate 3000 HPLC system (Thermo Scientific) and separated across a 4.6 mm × 50 mm ProSwift RP-1S column (Thermo Scientific). Peptides were eluted using a 1 ml/min gradient over 5 min from 1-35% Acetonitrile/0.1% TFA and fractions were collected every 30 seconds. Peptide fractions 1-15 were combined into odd and even fractions then dried.

### Mass spectrometric analysis

HPLC fractions were dissolved in loading solvent, and analysed by an Ultimate 3000 HPLC system coupled to a high field Q-Exactive (HFX) Orbitrap mass spectrometer (Thermo Scientific). Peptides were initially trapped in loading solvent, before RP separation with a 60 min linear acetonitrile in water gradient of 2-25 or 35% (HLA-I/HLA-II and trypsin analyses) across a 75 µm × 50 cm PepMap RSLC C18 EasySpray column (Thermo Scientific) at a flow rate of 250 nl/min. Gradient solvents contained additional 1%(v/v) DMSO and 0.1%(v/v) formic acid. An EasySpray source was used to ionise peptides at 2000 V, which were analysed by data-dependent acquisition. Initially a full-MS1 scan (120,000 resolution, 60 ms accumulation time, AGC 3×10^6^) was followed by 20 data-dependent MS2 scans (60,000 resolution, 120 ms accumulation time, AGC 5×10^5^), with an isolation width of 1.6 m/z and normalised HCD energy of 25%. For HLA-II charge states of 2–4 were selected for fragmentation, for HLA-I 1+ ions were also included. Dynamic exclusion was set for 30 s. For enzymatic digests of S protein normalised HCD was increased to 28%, only 2-4 charge states were acquired.

### Mass spectrometry peptide data analysis

Raw data files were analysed in PEAKS X software (Bioinformatic Solutions) using a protein sequence fasta file containing 20,606 reviewed human Uniprot entries downloaded on 24/05/2018, supplemented with the sequence for SARS-CoV-2 full-length S protein cloned. No enzyme specificity was set, peptide mass error tolerances were set at 5 ppm for precursors and 0.03 Da for MS2 fragments. Additionally, post translational modifications were identified utilising the Peaks PTM inbuilt *de novo*-led search for 303 common modifications. A 1% false discovery rate (FDR) was calculated using decoy database search built into Peak data plotting. Data manipulation was performed in R and excel. NetMHCII pan 4.0^23^ (http://www.cbs.dtu.dk/services/) was installed locally and utilised to define binding (rank score cut of 10). Sequence logos were generated by Seq2logo2.0. Venn diagrams and UpsetR plots were created using the online http://www.interactivenn.net/index2.html^61^ and UpsetR program in R. Clustering of sequences was done by GibbsCluster2.0 with defaults for MHC class II ligands and 1–5 clusters, the most obvious 1-2 clusters were selected. Peptides were aligned into nested sets using Immune Epitope Database (IEDB) Epitope Cluster Analysis^62^ tool http://tools.iedb.org/cluster/ and positions in proteins were determined by http://tools.iedb.org/immunomebrowser/. A python script was written to calculate the frequency of amino acids across the full length S protein. Jalview 2.11.1.0 ^63^ was used for multisequence alignment with MAFFT (Multiple Alignment using Fast Fourier Transform) with default settings. Alignments were visualised in Jalview and colouring amino acids by property based on CLustal_X definitions or identity. All graphs were plotted in R or Excel.

### Glyco-site and glycopeptide analysis

Raw data for the de-*N*-glycosylated samples were analysed as described above with the following modifications. A standard PeaksDB search was utilised with additional variable post translational modifications set for ^18^O labelling (C-terminal +2.00 Da), ^18^O deamidation (NQ, +2.99 Da), deamidation (NQ, −0.98 Da) and oxidation (M, +15.99 Da) and a fixed modification for carboxyamidomethylation (C, +57.02). Each site was checked manually for over-lapping peptides with consistent heavy deamidation present. Glycopeptides were identified by Byonic v3.8.13 (Protein Metrics, CA, USA). Spectra were extracted and searched against the same fasta file as above, but with unspecific cleavage sites and allowing unlimited missed cleavages. A library of 132 possible *N*-glycan compositions was utilized in a ‘rare’ variable modification search strategy. A fixed modification of carboxyamidomethylation was included for digested S. Data was filtered at a 1% protein FDR resulting in a 1.38 % peptide FDR in the combined dataset, glycopeptides were further filtered to ensure that oxonium ions were present in the MS2 spectrum and that scores were above 150 as described^64^. All reported glycopeptides were manually checked. Filtering and spectral counting was performed in Byologic v3.8.11, spectra with the same precursor mass and peptide identity was grouped and compositions determined as per the library.

## ACKNOWLEDGEMENTS

This study was funded by a CRUK Accelerator Award (C328/ A21998) to RP and NT; and NIH, NIAID, DAIDS Consortium for HIV/AIDS Vaccine Development (CHAVD) grant UM1 AI 144371 (KS, BFH, PA and PB). PB is a Jenner Institute Investigator. MTA was supported by a Macquarie University Safety Net grant. RK was funded via an Early Career Research Fellowship from the Cancer Institute NSW. We like to thank Paul Klenerman for his support and encouragement to carry out this study.

## AUTHOR CONTRIBUTIONS

RP, TP, PB and NT designed experiments, RP and TP conducted experiments. RP, TP, CW, MA established laboratory methodologies. RP, TP, RK, JP,MTA analysed the data. VS, RPD, YAM, EL, KS, BFH and PA provided reagents for this study. RP, TP, CW,PB, NT wrote the manuscript. PB and NT supervised this study and acquired funding. All authors contributed to revision of the manuscript.

