## Supplemental Figures for "Mapping the SARS-CoV-2 spike glycoprotein-derived peptidome presented by HLA class II on dendritic cells"

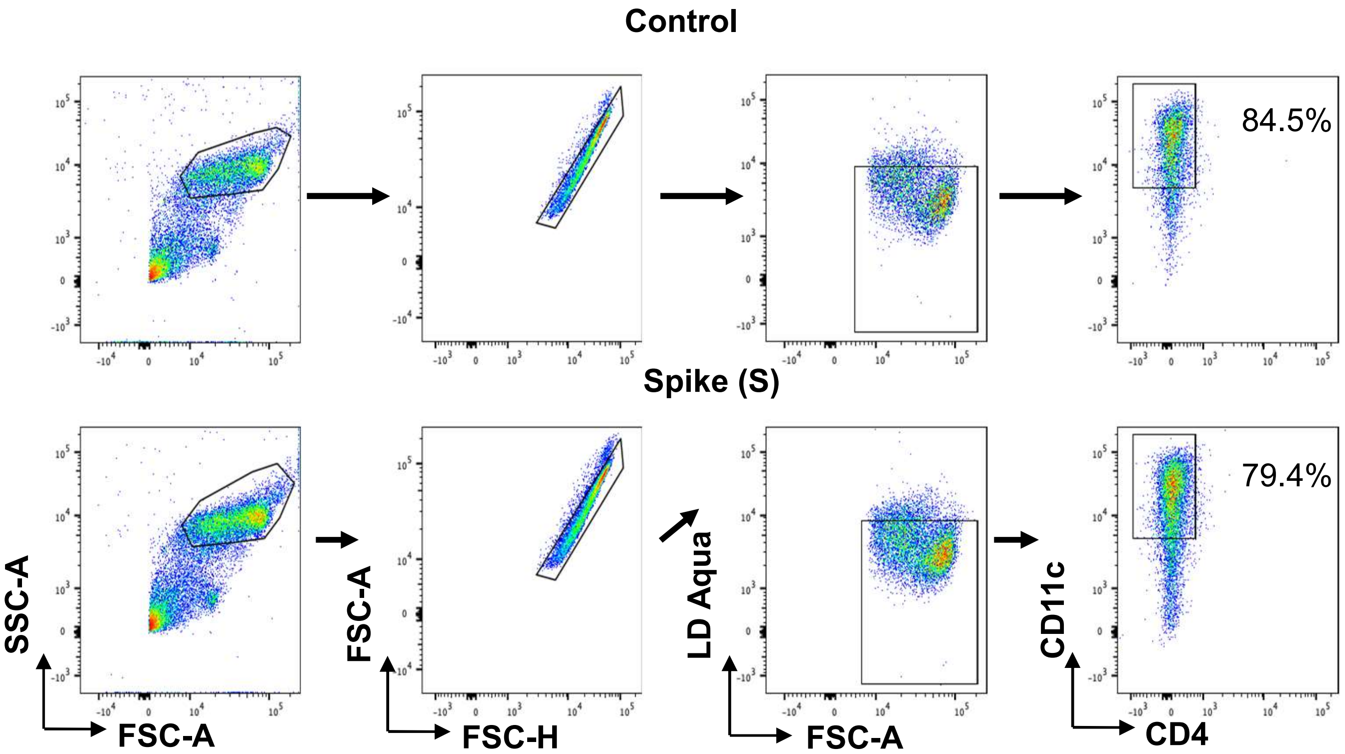


**Figure S1. Gating strategy for flow cytometry analysis of MDDCs**. The dotplots show examples of the gates applied to identify MDDCs in samples from one representative donor after pulsing with S (top) or control protein (bottom). Serial gates were applied to identify large mononuclear cells, singlets, live cells and CD11c+ MDDCs (the percentage of which within the live mononuclear cell population is indicated).

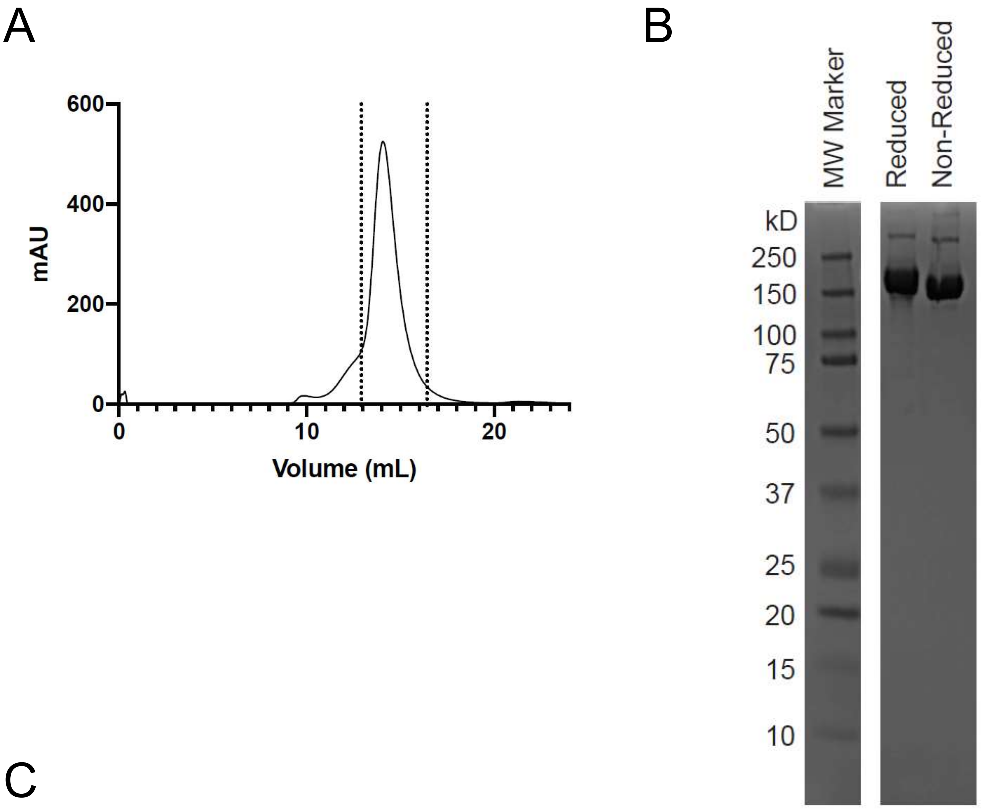


C

Furin cleavage site

SPIKE_SARS2 651 IGAEHVNNSYECDIPIGAGICASYQTQTNSPRRARSVASQSIIAYTMSLG
Recombinant SPIKE_SARS2* 651 IGAEHVNNSYECDIPIGAGICASYQTQTNSPGSASSVASQSIIAYTMSLG

SPIKE_SARS2 951 VVNQNAQALNTLVKQLSSNFGAISSVLNDILSRLDKVEAEVQIDRLITGR
Recombinant SPIKE_SARS2* 951 VVNQNAQALNTLVKQLSSNFGAISSVLNDILSRLDPPEAEVQIDRLITGR

**Figure S2. Production, purification, and sequence variation of the SARS-CoV-2 S ectodomain.** A. Size exclusion chromatography profile of the SARS-CoV-2 S protein that was purified using the C-terminal TwinStrep tags. The S protein was run on a Superose 6 Increase 10/300 column. The dotted lines indicate the portion of the peak that was collected and used for this study. B. SDS-PAGE gel of the purified S protein with lanes from left to right showing molecular weight marker, S protein run under reducing conditions and S protein run under non-reducing conditions. C. Pairwise sequence alignment of regions of the S protein sequence (P0DTC2, SPIKE_SARS2, displayed as top sequence) and corresponding regions of the sequence of the recombinant S protein employed in this study (bottom sequence). Peptides found in the immunopeptidome are indicated in red text, and the sites where the amino acid sequence of the recombinant protein differs from the Uniprot S sequence are highlighted in yellow.

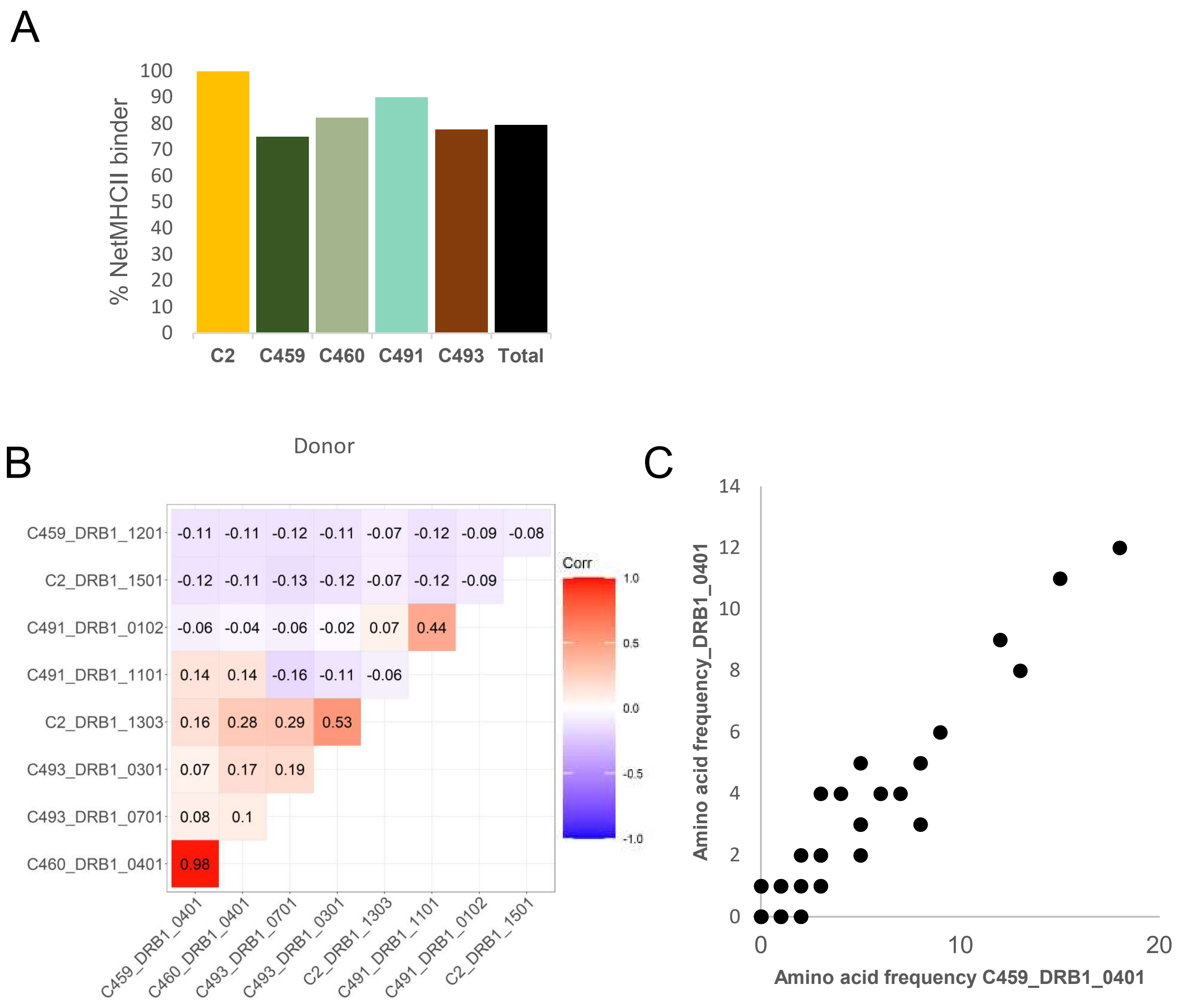


**Figure S3. Binding prediction and sequence correlation for peptides identified in the HLA-DR pulldown.** A. Proportion of HLA-DR peptide sequences predicted to bind to each donor's HLA-DR alleles by NetMHCIIpan 4.0. B. Correlation matrix indicating the concordance in the sequence of the peptides predicted to bind to the HLA-DR alleles indicated in each donor. Pearson correlation coefficients of the frequencies with which each amino acid in the S protein is represented in the peptides predicted to bind the indicated donor DR alleles and R squared values are shown; the direction and magnitude of the correlation is indicated by colour. C. Scatter plot of the frequency (number of times) with which each individual amino acid position within S is represented in all eluted S peptides predicted to bind to HLA-DRB1*04:01 in the two donors who shared this allele. Residues within S which were not embodied by peptides predicted to bind HLA-DRB1*04:01 are not plotted.

A

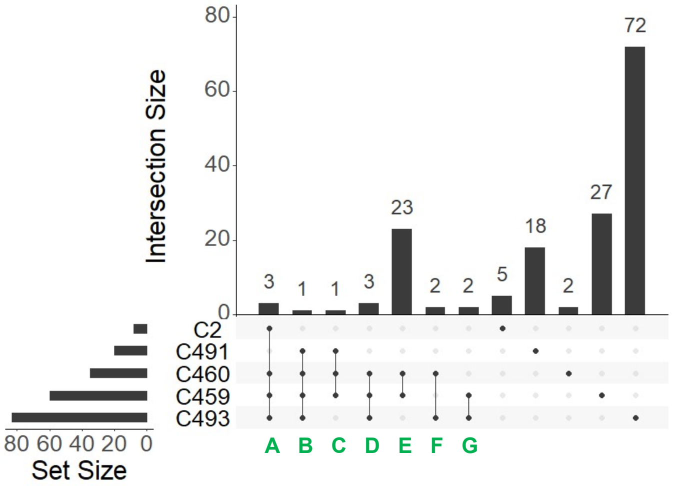


B

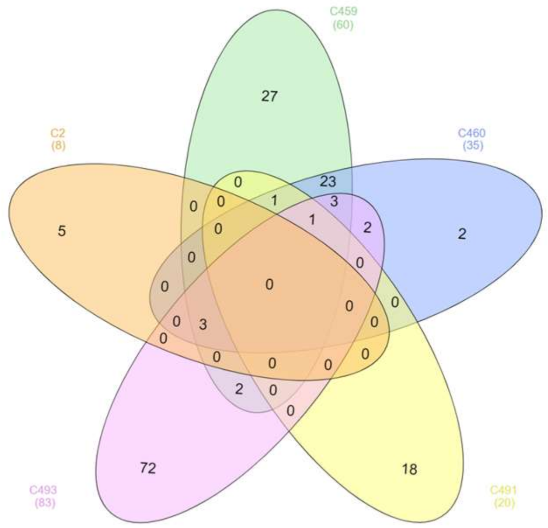


C

**A**

**C2 and C459 and C460 and C493:**
RGVYYPDK
SNLKPFERDISTEIYQA
SNLKPFERDISTEIYQ-

**B**

**C459 and C460 and C491 and C493:**
SNLKPFERDISTEIYQAG

**C**

**C459 and C460 and C491:**
RGVYYPDKVFRSSVL

**D**

**C459 and C460 and C493:**
RGVYYPDKVFRS
RGVYYPDKVFR-
LKPFERDISTEIYQ

**Figure S4. Peptide overlap between analysed donors.** A. UpSet R plot showing the number and degree of co-occurrence of the same S peptide sequences in the HLA-DR-bound immunopeptidomes of different donors. B. Venn diagram representing the data from (A), demonstrating that there were no S peptides that were detected in the HLA-II-bound immunopeptidomes of all donors. C. Sequences of the S peptides found to be presented in more than 3 donors.

A

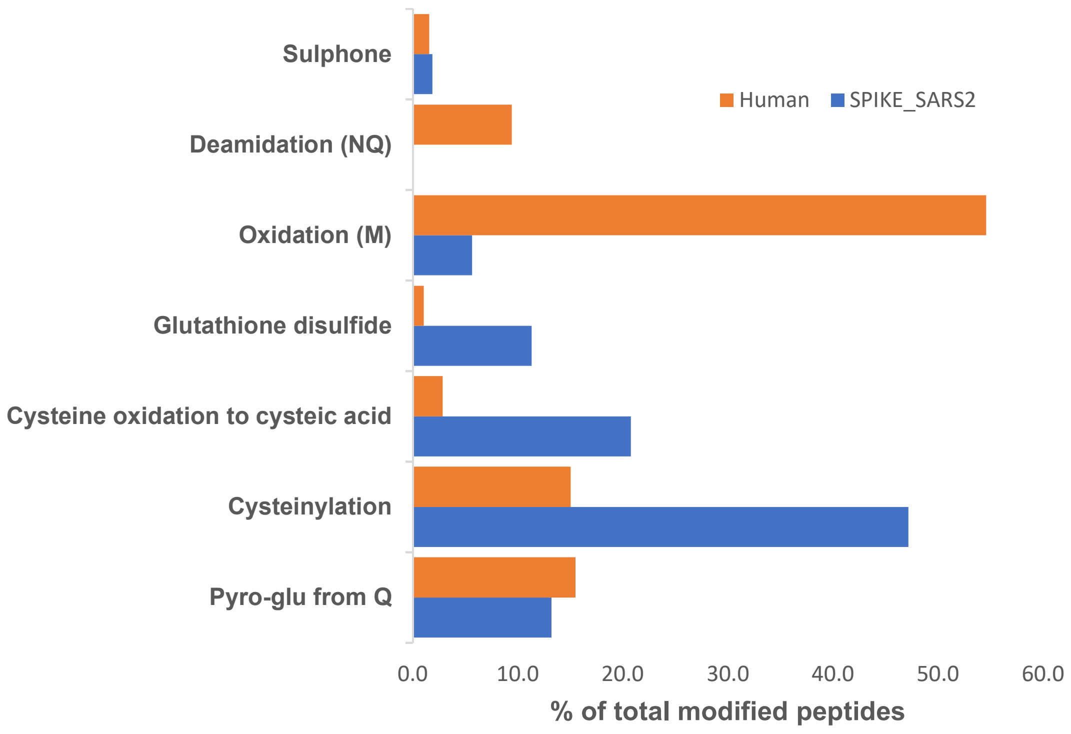


B

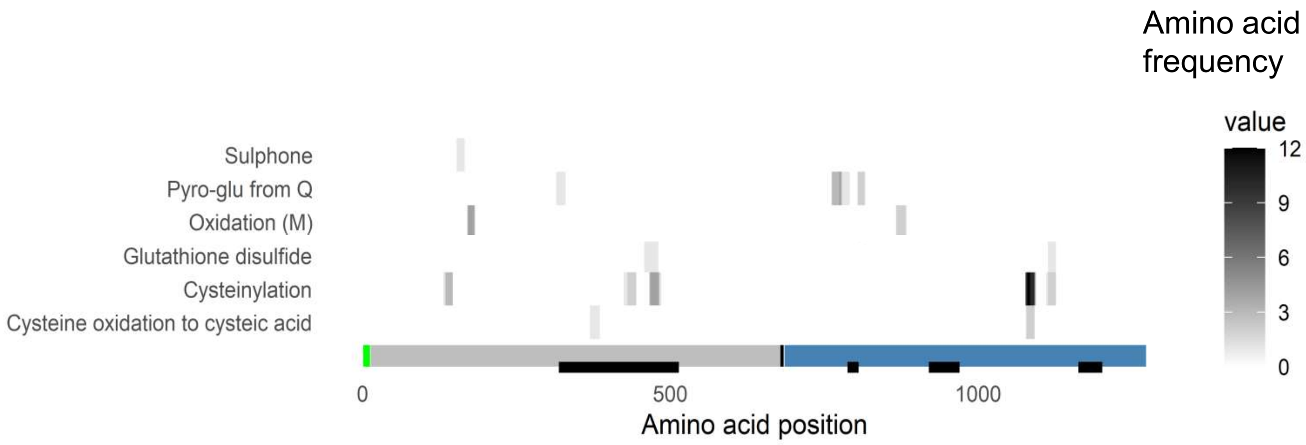


**Figure S5. PTM profile of HLA-II peptides.** A. Bar graph showing the most common peptide modifications detected as a proportion of all modifications reported (A-score >500), in both peptides mapping to S (blue) or mapping to human proteins (orange). B. Heatmap of positional amino acid frequency in S for the most commonly modified S peptide sequences identified in the HLA-II-bound immunopeptidome.
